# Multi-layer colocalization prioritizes adipose RNA regulatory features beyond gene expression at type 2 diabetes loci

**DOI:** 10.64898/2026.11.23.768771

**Authors:** Yuelong Chen, Shiya Huang, Tengxiao Lei, Yanci Wang, Haokun Han, Jiawei Li, Lixin Cheng, Weidong Tang, Jijun Tang, Nana Jin

## Abstract

Type 2 diabetes (T2D) genetic signals reside predominantly in non-coding regions, and linking them to molecular mechanisms requires resolving the RNA phenotypes through which they act in disease-relevant tissues. Adipose tissue is central to systemic glucose homeostasis, yet most colocalization studies have relied on total gene expression, which captures only one dimension of RNA regulation. Here, we integrated 338 fine-mapped T2D signals from DIAMANTE with five GTEx RNA measurements—gene expression, exon expression, transcript usage, transcriptional-event usage, and splice-junction usage—across subcutaneous and visceral adipose tissue from the eQTL Catalogue. Enforcing bilateral posterior coverage and a cumulative posterior selection rule across 13,216 admissible comparisons expanded testable T2D signals from 253 with gene expression alone to 293 and identified 77 colocalized pairs defining 38 signal–gene relationships across 30 T2D signals, with 87.0% of retained pairs involving non-gene-expression features. At loci with direct same-gene comparators, including BDNF-AS and PGM1, specific RNA features strongly colocalized with T2D while bulk gene expression favored distinct causal variants, demonstrating that gene-level analysis alone can miss or misattribute the primary regulatory signal. Locus-resolved fine-mapping linked a BDNF-AS splice junction to rs6265 in both depots and narrowed PGM1 and RCCD1 to shared or nested candidate-variant sets around defined transcript anchors. External adipose cohorts provided complementary support, single-nucleus and cell-class eQTL data defined cellular contexts, and metabolic-trait and multi-tissue analyses distinguished insulin-resistance (IRS1), fat-distribution (CCDC92), and cross-tissue adiposity (BDNF-AS) profiles. Together, these results provide a multi-layer adipose regulatory atlas that converts T2D signals into specific RNA features, candidate variants, cellular settings, and measurement-led routes for functional validation.

**Research in context:** *What is already known about this subject?:* Most type 2 diabetes (T2D) genetic risk signals lie in non-coding regions and act through gene regulation in metabolic tissues such as subcutaneous (SAT) and visceral (VAT) adipose tissue. Previous adipose colocalization studies have relied predominantly on bulk gene expression, which captures total transcript abundance but can miss genetic effects acting on isoform usage, exon inclusion, alternative promoters or RNA splicing.

*What is the key question?:* How do transcriptomic measurements beyond total gene expression expand the testable T2D signal space in SAT and VAT, and how does feature-level colocalization compare with same-gene bulk expression at T2D susceptibility loci?

*What are the new findings?:* Integrating 338 fine-mapped T2D signals with five RNA layers across SAT and VAT expanded testable signals from 253 to 293 and identified 77 colocalized pairs defining 38 signal–gene relationships across 30 T2D signals, with 87.0% of retained pairs involving non-gene-expression features. At loci including BDNF-AS (converging on rs6265) and PGM1, specific splice junctions or transcripts strongly colocalized with T2D whereas bulk gene expression favored distinct causal variants, a dichotomy corroborated across external adipose cohorts (e.g., USP36 splicing vs WFS1 expression in METSIM). Combining single-nucleus and cell-class eQTLs with cardiometabolic traits distinguished adipocyte insulin-resistance (IRS1), adipose-restricted fat-distribution (CCDC92) and brain–adipose adiposity (BDNF-AS) profiles, specifying exact RNA readouts and cellular settings for follow-up.

*How might this impact on clinical practice in the foreseeable future?:* By shifting target prioritization from broad gene nomination to specific RNA isoforms, splice junctions and cellular contexts alongside curated bioactivity profiles, this atlas provides an actionable roadmap for developing and testing isoform-resolved therapeutic strategies in T2D.

## Introduction

Genome-wide association studies and trans-ancestry fine-mapping consortia such as DIAMANTE have defined hundreds of independent type 2 diabetes (T2D) signals^1,2^, the vast majority of which reside in non-coding regions^3^. Translating these signals into biological mechanisms requires linking them to molecular phenotypes in disease-relevant tissues. White adipose tissue is central to systemic glucose homeostasis, insulin sensitivity, and lipid storage^4-6^, and subcutaneous (SAT) and visceral adipose tissue (VAT) differ in developmental origin, cellular composition, and metabolic function^7-9^. Mapping how T2D signals regulate molecular phenotypes across both depots is therefore essential for understanding the peripheral mechanisms of T2D susceptibility.

Colocalization between disease signals and molecular quantitative trait loci (QTLs) provides a direct framework for this mapping^10,11^. Gene expression (GE) has been the default readout, and GE colocalization has nominated many candidate genes at metabolic loci^12-14^. Total transcript abundance, however, captures only one dimension of RNA regulation. Genetic effects on promoter choice, exon inclusion, transcript usage, and splice-junction usage can alter RNA structure^15-17^ or isoform composition without changing overall abundance. In adipose tissue, splice-junction QTLs have already implicated RNA features at cardiometabolic signals^18^, including features without a detected gene-level eQTL. These findings indicate that the molecular interpretation of a T2D signal can depend on which RNA phenotype is measured. Uniformly processed resources such as the eQTL Catalogue, which quantifies five RNA layers ^19^within a harmonized framework, now make it possible to evaluate this systematically across tissues and depots.

T2D colocalization studies have largely focused on GE, often within a single depot or at selected loci. Two questions remain open: how non-GE features expand the testable signal space, and how feature-level evidence compares with same-gene GE evidence at signals where both are evaluable. Addressing these questions requires an analytical framework that enforces bilateral posterior coverage and resolves competing signals across layers and depots, together with a biological framework that connects retained features to external adipose cohorts, cellular contexts, and physiological traits.

Here, we integrated 338 fine-mapped T2D signals from DIAMANTE^2,19,20^ with five RNA measurements—GE, exon expression, transcript usage, transcriptional-event usage, and splice-junction usage—in SAT and VAT, using GTEx QTLs uniformly processed by the eQTL Catalogue. From 50,995 candidate signal–component pairs, 13,216 met a bilateral shared-variant posterior coverage requirement of at least 0.80 together with component and variant integrity checks. A cumulative mean-score selection rule retained 77 component pairs, defining 38 signal–gene relationships across 30 T2D signals. Multi-layer integration expanded the number of testable signals from 253 with GE alone to 293, and 67 of the 77 retained pairs involved non-GE measurements. At loci with direct same-gene comparators, such as BDNF-AS and PGM1, specific RNA features strongly colocalized with T2D while the evaluated GE component favored distinct causal variants (PP.H3 > 0.999), showing that locus interpretation depends on the measured RNA layer.

Locus-resolved fine-mapping of BDNF-AS, PGM1, and RCCD1 then sharpened these observations into concrete molecular hypotheses. At BDNF-AS, usage of a single splice junction (chr11:27,640,005– 27,659,171) colocalized with T2D in both depots and converged on rs6265 as the leading candidate variant conditional on H4, whereas VAT BDNF-AS expression favored distinct variants. At PGM1 and RCCD1, transcript-usage and transcriptional-event measurements narrowed candidate variant sets around shared or nested transcript anchors. Evaluation across four external adipose cohorts (FUSION, TwinsUK, METSIM, and KOBS) provided complementary support for prioritized relationships, including the locus-specific distinction between junction-driven and expression-driven signals. Single-nucleus expression and cell-class eQTLs placed these relationships into defined cellular environments, and metabolic-trait genetics, BMI-adjusted T2D effects, and multi-tissue QTLs defined their physiological profiles. Combining these findings with pharmacological annotations defined a measurement-led roadmap for targeted functional validation of prioritized T2D loci.

## Methods

### Study design and data resources

We integrated fine-mapped type 2 diabetes (T2D) signals with five molecular quantitative trait locus (QTL) layers in subcutaneous (SAT) and visceral (VAT) adipose tissue. We then characterized the retained relationships using transcript structure, external adipose cohorts, single-nucleus expression, cell-class expression QTLs, metabolic-trait genetics, tissue context and pharmacological annotations. The discovery unit was a disease signal–molecular component pair. Downstream summaries grouped retained pairs into signal–gene relationships. External, cellular, physiological and pharmacological analyses were applied to this fixed discovery set and did not change its membership. All inputs were publicly available summary statistics or de-identified data; no participants were recruited and no biological material was collected for this study.

Disease evidence came from the DIAMANTE 2022 fine-mapping archive, comprising 338 distinct signals at 237 loci ^2^. Molecular evidence came from the uniformly processed GTEx study in eQTL Catalogue release 7, with 581 SAT and 469 VAT donors ^19,20^. Ten datasets represented gene expression (GE), exon expression, transcript usage, transcriptional-event usage and splice-junction usage in each depot. Transcript usage was quantified relative to the other transcripts of the same gene.

Transcriptional-event usage was quantified with txrevise as the usage of promoter (upstream), internal (contained) or 3′-end (downstream) events relative to events of the same class within a gene ^16^. Splice-junction usage was quantified with LeafCutter as the usage of a junction within its intron cluster ^48^. The primary molecular evidence was the released per-variant log Bayes factors (BFs) of each SuSiE credible-set component, rather than marginal association statistics ^22^.

### Evidence transformation and shared-variant coverage

For disease signal *s* and variant *v*, the released base-10 log BF was converted to natural-log evidence, 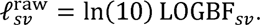 Annotation-informed disease BFs (BF*) were constructed over the complete released signal universe *U_G_*(*s*):

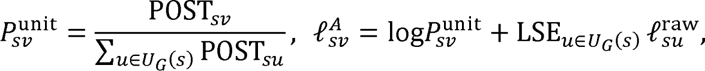

where POST is the released variant posterior and LSE the logarithm of the sum of exponentials. This transformation preserves the summed raw BF of each signal while incorporating the published annotation-informed posterior allocation. Released posterior zeros remained zero BF contributions, and disease-side variant priors were uniform because annotation information was already included in BF*. Molecular component posteriors were reconstructed from the released natural-log BFs over each complete component universe 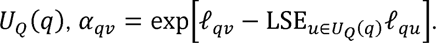

GRCh37 disease variants were mapped to GRCh38 with pinned UCSC chain files ^23^, requiring a unique reciprocal mapping and agreement with the reference allele. Shared variant sets contained one-to-one coordinate and allele matches. Molecular indels remained in the complete-component denominators but could not match the biallelic disease release. Uniquely represented palindromic single-nucleotide variants were retained in the primary analysis and removed in a sensitivity analysis.

For the shared set *S_j_* of pair *j*, disease coverage was *C_G, j_* = ∑_*v*∈*S_j_*_ POST_*sv*_ and molecular coverage was *C_Q, j_* = ∑_*v*∈*S_j_*_ *α*_*sv*_. Disease coverage used the released posterior before unitization, and neither coverage was renormalized to the intersection. A pair was admissible when both unrounded coverages were at least 0.80 (bilateral coverage) and component and variant identities were valid. Four pairs whose precision-propagated coverage intervals crossed 0.80 were excluded as unresolved boundary cases. A signal was testable in a dataset or layer when it had at least one admissible pair. Of 50,995 candidate pairs, 13,216 were admissible (Fig. 1b; Supplementary Figs. S1 and S2; Table S2).

**Figure 1.**
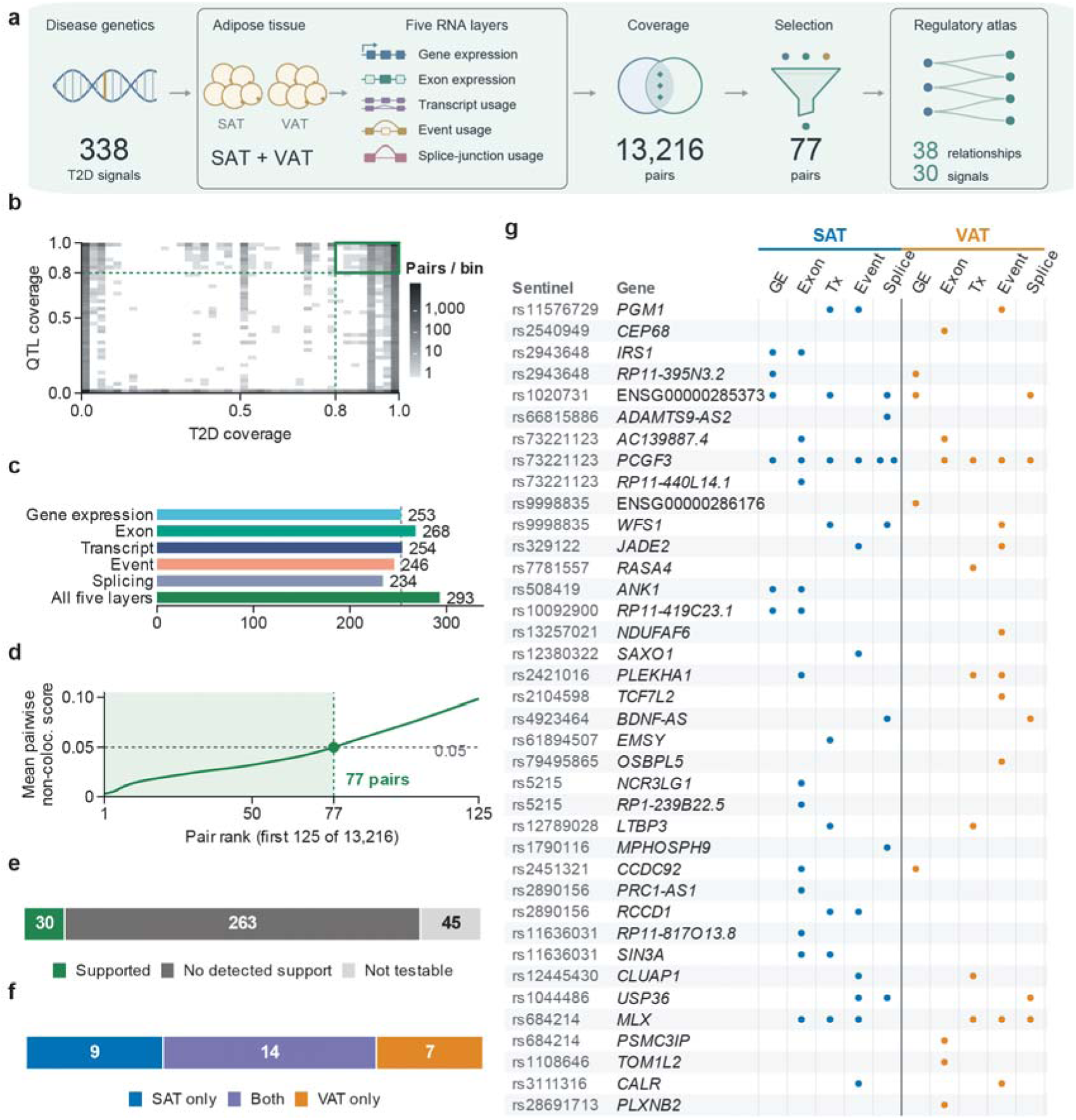
Multi-layer integration defines an adipose regulatory atlas of type 2 diabetes. (a), Integration of 338 fine-mapped type 2 diabetes (T2D) signals with five RNA measurements in subcutaneous (SAT) and visceral (VAT) adipose tissue. (b), Shared posterior coverage for 50,995 candidate signal–molecular component pairs. Shading shows counts per two-dimensional bin on a log scale. Green lines mark the bilateral 0.80 threshold; four source-precision boundary cases were excluded, leaving 13,216 admissible comparisons. (c), Signals with at least one admissible comparison in each molecular layer or their union. Integration adds 40 testable signals beyond the 253 testable with gene expression alone; these counts describe testability. (d), Selection boundary among all 13,216 ranked comparisons (first 125 shown). The largest complete-tie prefix with mean pairwise non-colocalization score ≤0.05 contains 77 pairs (mean 0.04985). (e), Complete signal outcomes: 30 supported, 263 tested without detected support and 45 not testable. (f), Dataset-level support among the 30 supported signals: nine in SAT alone, 14 in both depots and seven in VAT alone. Single-depot detections had testable comparisons in the other depot. (g), All 38 signal–gene relationships across ten depot–layer combinations. Each point represents one of the 77 retained component pairs; multiple pairs within a cell are offset. Rows show the disease sentinel and mapped gene, ordered by chromosome and locus. Blank cells indicate no retained pair. GE, gene expression; Tx, transcript usage; Event, transcriptional event usage; Splice, splice-junction usage.

### Colocalization model

Colocalization used coloc.bf_bf in coloc 6.0.1 with posterior trimming disabled ^10,11^. Per-variant priors were *p*_1_ = *p*_2_ = 10^-4^ and *p*_12_ = 5 × 10^-6^. For *Q* shared variants, a prior *p_t_* was replaced by 1/(*Q* + 1) when *Qp_t_* ≥ 1 and otherwise retained; these effective priors, *p̃_t_*, were passed explicitly. With 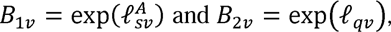 the hypothesis weights were 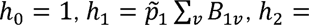 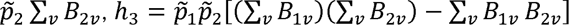 and 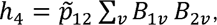 and normalization gave the posterior probabilities PP.H0–PP.H4. H0 denotes no association with either trait, H1 and H2 association with the disease or molecular trait only, H3 distinct causal variants and H4 a shared causal variant. Computation used log-space arithmetic. An independent implementation reproduced the posterior probabilities and conditional shared-variant weights to an absolute tolerance of 10 (Supplementary Fig. S1).

### Selection, attribution and signal classification

All 13,216 admissible comparisons formed a single selection family. Pairs were ordered by their non-colocalization score, *l_j_* = PP. H0_j_ + PP. H1_j_ + PP. H2_j_ + PP. H3_j_, and the selected set was the largest prefix ending at a complete block of exact ties with a mean score of at most 0.05 (Fig. 1d). Because *l_j_* is the model posterior probability that pair *j* does not colocalize, the mean score of the selected set is its model-based expected proportion of non-colocalized pairs, as in the direct posterior probability approach to false-discovery control ^24^. This expectation holds only to the extent that the colocalization posteriors are calibrated, which depends on the priors, the single-causal-variant model and linkage disequilibrium (LD). The rule therefore defined a prioritized discovery set and was not interpreted as an empirically calibrated false-discovery rate.

Attribution grouped comparisons sharing locus, depot, molecular layer, feature and molecular component. A pair was retained only when a single disease signal was selected within its group; groups with competing selected signals were classified as attribution-ambiguous (Supplementary Fig. S3d). Retained pairs were grouped by disease signal and gene into signal–gene relationships (Fig. 1g; Table S1). Each disease signal was classified, in order, as supported, attribution-ambiguous, tested without detected support or not testable (Fig. 1e). Selections under fixed thresholds of PP.H4 > 0.80 and PP.H4 > 0.90 were compared with the primary set using identical inputs (Supplementary Fig. S3a–c).

### Robustness and prioritization analyses

Complete sensitivity reruns repeated selection and attribution under joint priors *p*_12_ ∈ {5 × 10^-7^, 10^-6^, 2 × 10^-6^, 10^-5^}, after palindrome exclusion, with raw rather than annotation-informed disease BFs, and in a released-zero stress test that assigned posterior zeros a value of 10^-18^ before unitization. All reruns used bilateral coverage of 0.80 (Table S4).

To prioritize relationships for detailed follow-up, the complete analysis was additionally repeated at the primary prior with bilateral coverage requirements of 0.90 and 0.95. These reruns identify relationships whose shared-variant evidence is most completely represented in both datasets; they were used only as follow-up priority tiers and did not define alternative result sets. For each of the 38 relationships, we recorded the lowest tested joint prior at which it remained retained (descending from the primary 5 × 10^-6^ through 2 × 10^-6^, 10^-6^ and 5 × 10^-7^) and the highest coverage tier at which it remained retained (0.80, 0.90 or 0.95). Retention was nested along both series for every relationship, so these two positions were well defined. They were displayed jointly but derive from separate reruns and do not constitute a joint parameter test (Fig. 2e). No composite priority score was calculated.

**Figure 2.**
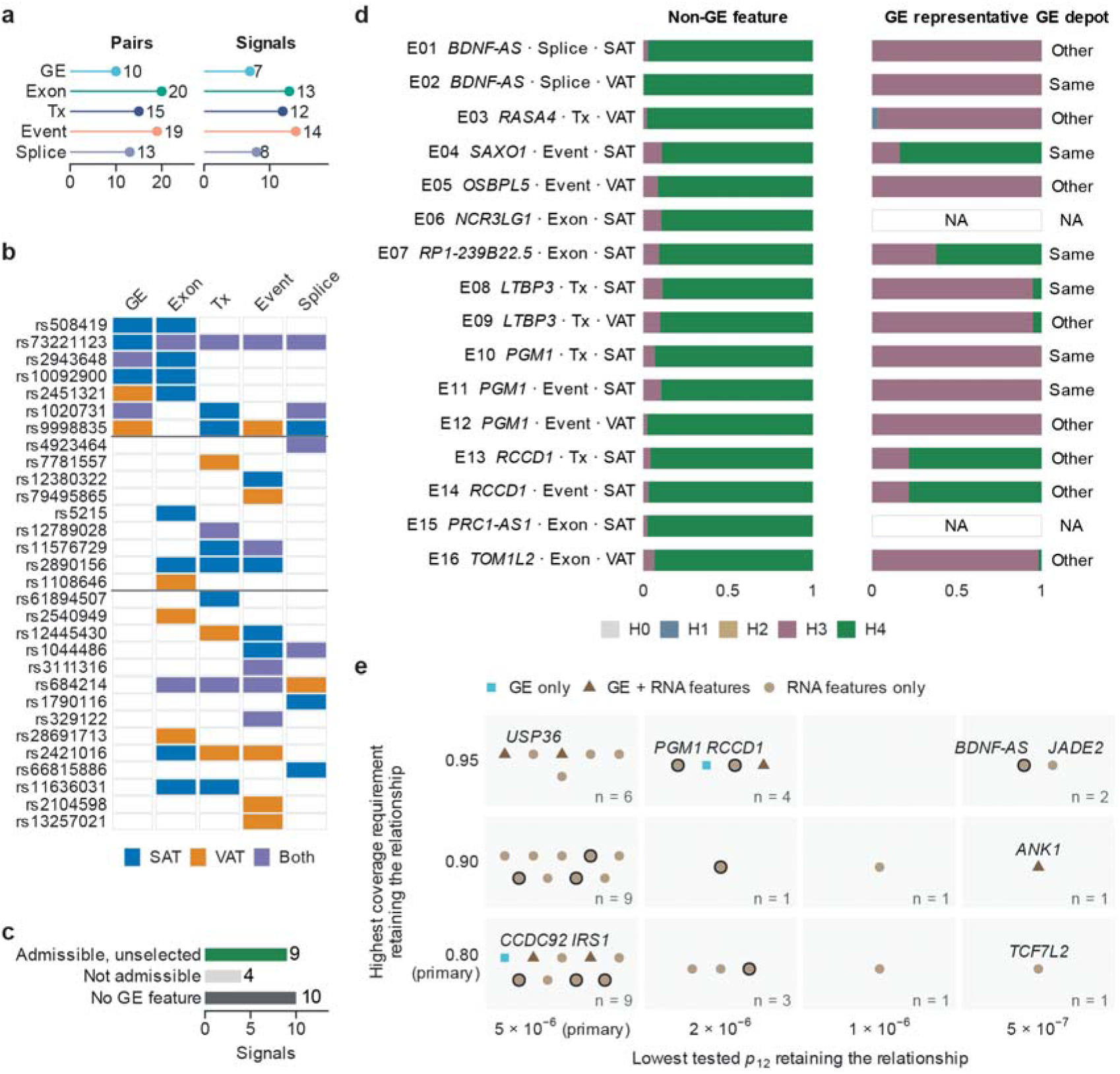
Molecular layers extend T2D signal interpretation beyond gene-level expression. (a) Retained pairs and supported signals by layer. (b) Layer/depot support for all 30 supported signals, grouped by GE comparator status; blank cells indicate no retained pair. (c) GE comparator status among the 23 signals with a retained non-GE pair but no retained GE pair: admissible but unselected (9), not admissible (4) or no comparable GE feature (10), using either depot. (d) H0–H4 compositions for the 16 retained non-GE molecular-feature–depot pairs at the nine signals with an admissible but unselected GE comparison (rows: pair ID, gene, layer and discovery depot) and for each pair’s highest-H4 admissible same-gene GE comparison across either depot, with pair-ID tie-breaking. Exact feature identities, GE comparator keys and comparator status are given in Table 1, and complete comparator records in Table S4. These are different RNA measurements of the same gene, not two analyses of one feature. GE depot gives whether the displayed representative is from the same depot as the non-GE pair (six pairs; an admissible same-depot comparison exists for eight); NA denotes no admissible same-gene GE comparison (E06, no same-gene GE feature; E15, comparison failed the bilateral 0.80 coverage requirement). (e) Discovery-stage position of all 38 signal– gene relationships: the lowest tested joint colocalization prior (p12) at which each relationship remained retained, at bilateral coverage 0.80 (x), and the highest bilateral coverage requirement at which it remained retained, at the primary prior (y). The axes come from separate complete reruns of selection and attribution, not a joint parameter test; the 0.90 and 0.95 coverage reruns are follow-up priority tiers, not alternative result sets. All 38 relationships were also retained under the more permissive prior tested. Colour and shape give the retained molecular evidence; outlined points are the 11 relationships of d and Table 1. Labels mark all relationships retained at the lowest tested prior and representative relationships analysed in later sections; positions within a cell carry no meaning. Complete branch status is given in Table S4. GE, gene expression; Exon, exon expression; Tx, transcript usage; Event, transcriptional-event usage; Splice, splice-junction usage. RP1-239B22.5, ENSG00000260196.

Within-phenotype thinning tested whether results depended on retaining several components of the same molecular phenotype. Three rules retained one component per dataset, layer, gene and molecular-trait identity: the minimum component index, the maximum component index, or the component with the smallest SHA-256 hash of a fixed seed and its identifiers. Phenotypes whose identity could not be resolved remained unthinned, and selection and attribution were repeated on each reduced family (Supplementary Fig. S3e).

Released disease posteriors were recorded to 18 decimal places. Five precision perturbations varied each value within its rounding interval and within [0, 1], after which BF* reconstruction, colocalization, selection and attribution were repeated and compared with a zero-perturbation control (Table S3).

### Molecular features and gene-expression comparators

Layer and depot summaries were derived from the retained pairs (Fig. 2a,b). A signal was classified as non-GE-only when it had at least one retained non-GE pair and no retained GE pair. For each such signal, we classified the GE evidence for the implicated genes in either depot as admissible but unselected, inadmissible (coverage not met) or absent (no comparable GE feature) (Fig. 2c). At signals with admissible but unselected GE evidence, the GE representative of each retained non-GE pair was the admissible same-signal, same-gene GE pair with the highest PP.H4 in either depot, with ties resolved by pair identifier; whether the representative came from the same depot was recorded (Fig. 2d; Table S4).

The molecular features of all retained pairs were mapped to eQTL Catalogue phenotype metadata (v23.04.1) and GENCODE v39 transcript structures ^25^. Junction boundaries and transcript models were recorded separately from any inferred protein consequence. For locus-resolved displays, variant posteriors conditional on H4 were obtained by normalizing *B*_1*v*_*B*_2*v*_ over the shared variants, and 95% conditional candidate sets accumulated posterior mass in descending order while retaining complete boundary ties (Fig. 3; Supplementary Fig. S4; Table S5). These sets describe localization under the shared-variant model and were not treated as independent colocalization evidence.

**Figure 3.**
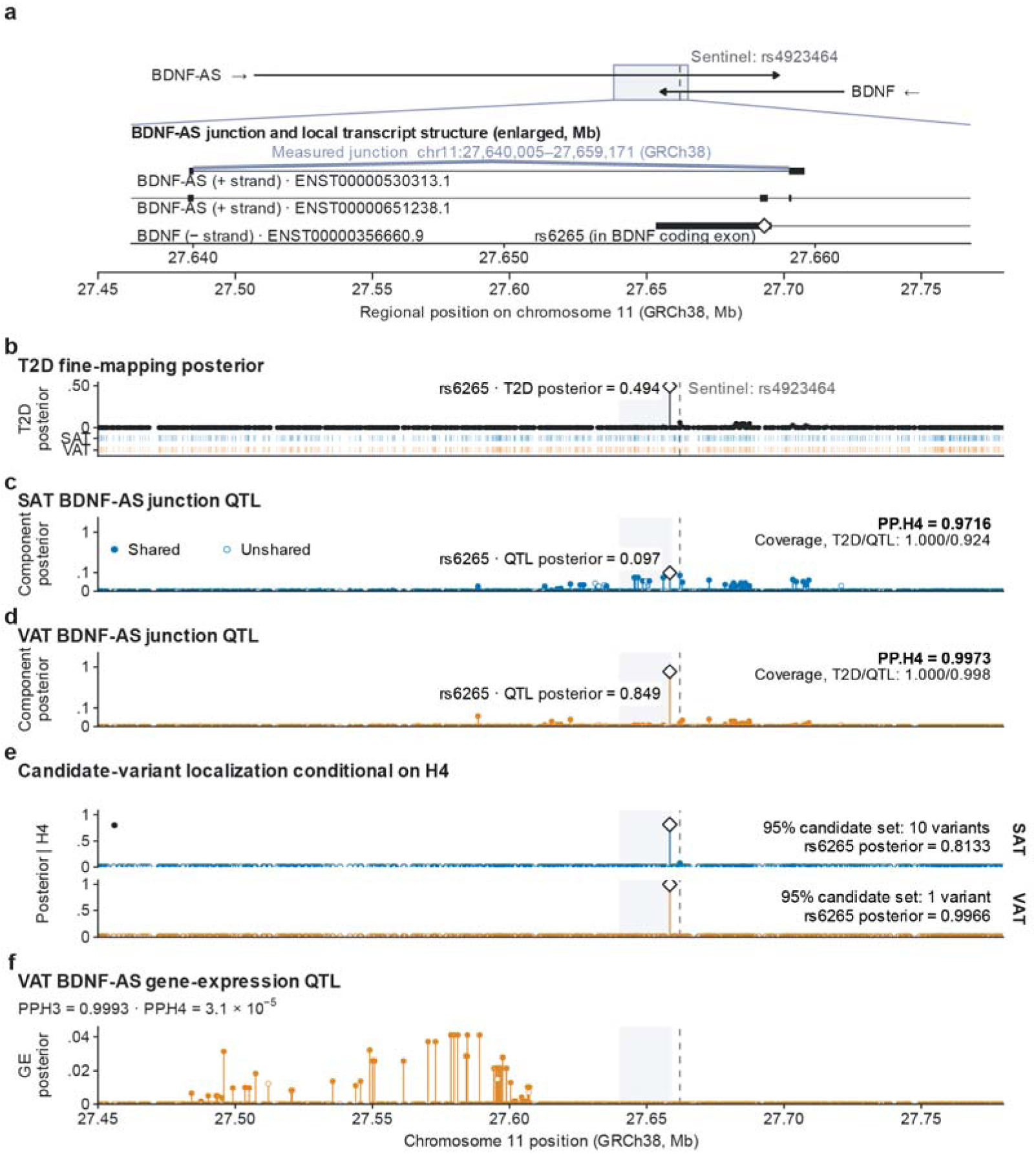
Locus-resolved colocalization prioritizes a BDNF-AS splice junction and rs6265 in both adipose depots. (a) BDNF/BDNF-AS region on the coordinates of b–f; the shaded box marks the interval enlarged below, which shows GENCODE v39 models of the junction-bearing BDNF-AS transcript, an alternative BDNF-AS transcript and BDNF. The arc marks the measured junction; junction usage does not identify a unique full-length isoform. rs6265 (diamond) lies inside the junction interval, in the BDNF coding exon. (b) Released T2D fine-mapping posterior; rs6265 is the fine-mapping lead and rs4923464 the reported sentinel. Rugs mark variants shared with the SAT and VAT junction components. (c,d) Complete-source posteriors of the SAT and VAT junction components (L1) on a common square-root scale; filled points are shared variants. (e) Variant posteriors conditional on H4 for each depot; filled points form the 95% conditional sets (complete sets, including boundary ties and the sentinel in the SAT set, in Table S5). Conditional sets describe localization under the shared-variant model and are not independent colocalization evidence. (f) VAT BDNF-AS gene-expression component (ENSG00000245573, L1; G01-V in Table 1 and Fig. 2d), same-depot for d and cross-depot for c. Diamonds mark rs6265 and dashed lines the reported sentinel rs4923464; shading marks the junction interval. Tracks share a 330-kb window retaining over 99.96% of each disease/QTL posterior without renormalization.

### External adipose cohorts

External component-level comparisons covered 46 SAT signal–feature targets in FUSION (271 donors) and TwinsUK (381 donors) from eQTL Catalogue release 7 ^19,26,27^. These analyses reused the original disease signals and applied the primary component-based colocalization, coverage, selection and attribution procedure. Exact feature identity, availability of released statistics and testability were recorded separately (Fig. 4a; Table S6).

**Figure 4.**
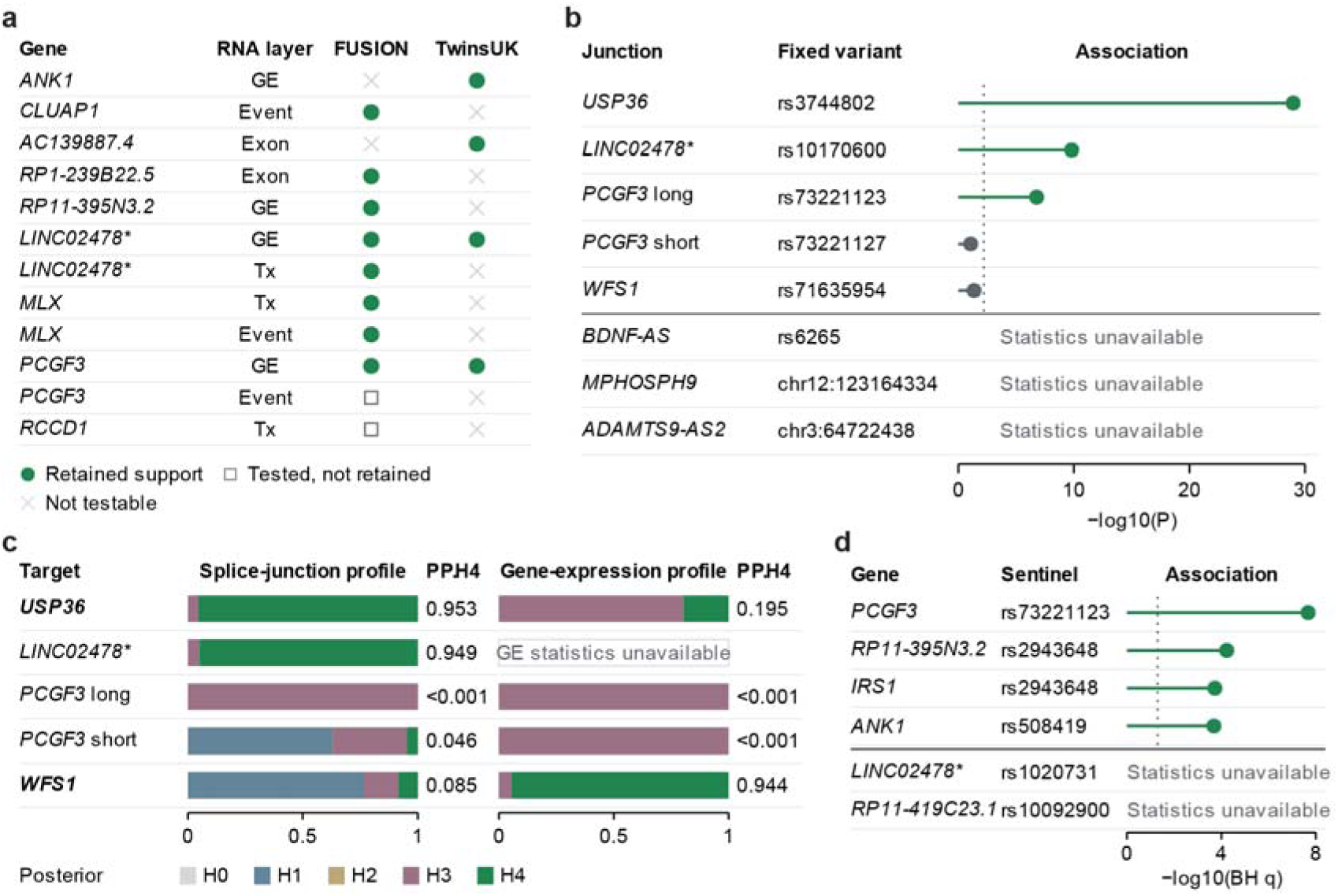
External adipose datasets provide complementary support for prioritized regulatory relationships. (a) Component-level colocalization in FUSION and TwinsUK for the 12 of 46 molecular targets with an admissible comparison in at least one cohort; the ten targets with retained support under the original selection and attribution procedure map to eight signal–gene relationships. (b) METSIM associations between discovery-fixed variants and the eight planned SAT splice-junction targets; dotted line, Bonferroni threshold for eight tests. Missing statistics are not negative evidence. Junction boundaries match, but phenotype equivalence (quantification denominator and strand) is unresolved. (c) METSIM regional H0–H4 compositions for the five junction targets with statistics, comparing the full released splice-junction profile with the gene-expression profile of the same gene under the primary model; all displayed profiles meet bilateral coverage 0.80. Low PP.H4 can reflect distinct variants (H3; USP36 gene expression, PCGF3) or no clear molecular association (H1; WFS1 junction). The USP36 and WFS1 contrasts persist on the common-variant domain (Table 2; parameter settings in Supplementary Fig. S5). A fixed-variant association (b) does not establish colocalization, as at the longer PCGF3 junction. (d) KOBS sentinel–gene associations for the six planned lookups; BH correction over six tests, dotted line q = 0.05; associations do not assign effect direction. Exact values are given in Table 2 and Supplementary Table S6. LINC02478*, discovery gene ENSG00000285373, labelled ITGB6 externally owing to transcript annotation reassignment; RP11-395N3.2, ENSG00000272622; RP11-419C23.1, ENSG00000254038; the USP36 junction (signal CYTH1.rs1044486) carries the external label CYTH1,USP36. GE, gene expression; Tx, transcript usage; Event, transcriptional-event usage.

METSIM provided marginal SAT association summaries from 426 participants ^18^. We evaluated the eight SAT splice-junction targets of the external target set. Candidate variants were fixed in advance from the retained SAT discovery pairs as all variants tied for the maximum H4-conditional posterior, giving eight planned variant–junction tests. Coordinates, alleles, gene annotation, strand and cluster membership were checked separately. Bonferroni correction used all eight planned tests, and unavailable statistics remained missing (Fig. 4b). Junction-boundary agreement supported variant–junction comparison, whereas equivalence of quantification denominators and strand was assessed separately and remained unresolved.

Regional models compared the full released METSIM splice-junction profile and the gene-expression profile of the same gene with the disease signal, and repeated both comparisons on the common disease–splice–expression variant domain. Two-sided P values were converted to *z* = < Φ^-1^ (1" *P*/2), the variance of the molecular effect was approximated as *V* = [2*Nf*(1"*f*)]^-1^ with *N* = 426 and source allele frequency *f*, and with *W* = SD^2^ and *r* = *W*/(*V* + *W*) the molecular log approximate BF was 1⁄2 [log (1-*r*) + *rz*^2^] ^28^. The primary model used disease BF*, a molecular-effect prior SD of 0.15 and *p*_12_ = 5 × 10^-6^, retained palindromic variants and applied no additional frequency filter (Fig. 4c). Coverage was calculated against the complete source profiles, and common-domain models required coverage of the disease, splice and GE profiles. Sensitivity branches varied the joint prior, molecular-effect prior, disease BF representation, palindrome exclusion and a minor-allele-frequency filter (Supplementary Fig. S5).

KOBS bulk SAT expression data were used to test the six original gene-expression relationships at their disease sentinels in 262 participants ^29^. Benjamini–Hochberg (BH) adjustment used all six planned tests ^30^; unavailable lookups were retained in the multiplicity family with a placeholder P value of 1 while their source statistics remained missing (Fig. 4d; Table S6). These associations were not used to assign effect direction.

### Single-nucleus expression

We analyzed the human Emont adipose single-nucleus RNA-seq count matrix (GEO GSE176171), comprising 137,684 nuclei from 24 biosamples and 13 donors; SAT was available from all 13 donors and VAT from ten of them ^9^. Biosamples were pooled within donor, depot and source cell type. Four inconsistent SAT donor labels were resolved by exact barcode joins to the source metadata, and repeated VAT samples from the same donor were pooled. Stable Ensembl identifiers mapped 35 of the 38 candidate genes. For each donor–depot–cell-type pseudobulk, candidate UMI counts were divided by the total counts across all 29,093 released features, scaled to counts per million and transformed as log□(CPM + 1). Pseudobulks required at least 20 nuclei, and ranked cell-type contexts required at least three eligible donors. Cell-type summaries weighted donors equally, and within-donor nuclear detection fractions were averaged across eligible donors. Absent features remained missing rather than zero. Leave-one-donor-out analyses removed each donor across its sampled depots and repeated the ranking of cell types (Fig. 5b,d; Supplementary Figs. S6 and S7; Table S7).

**Figure 5.**
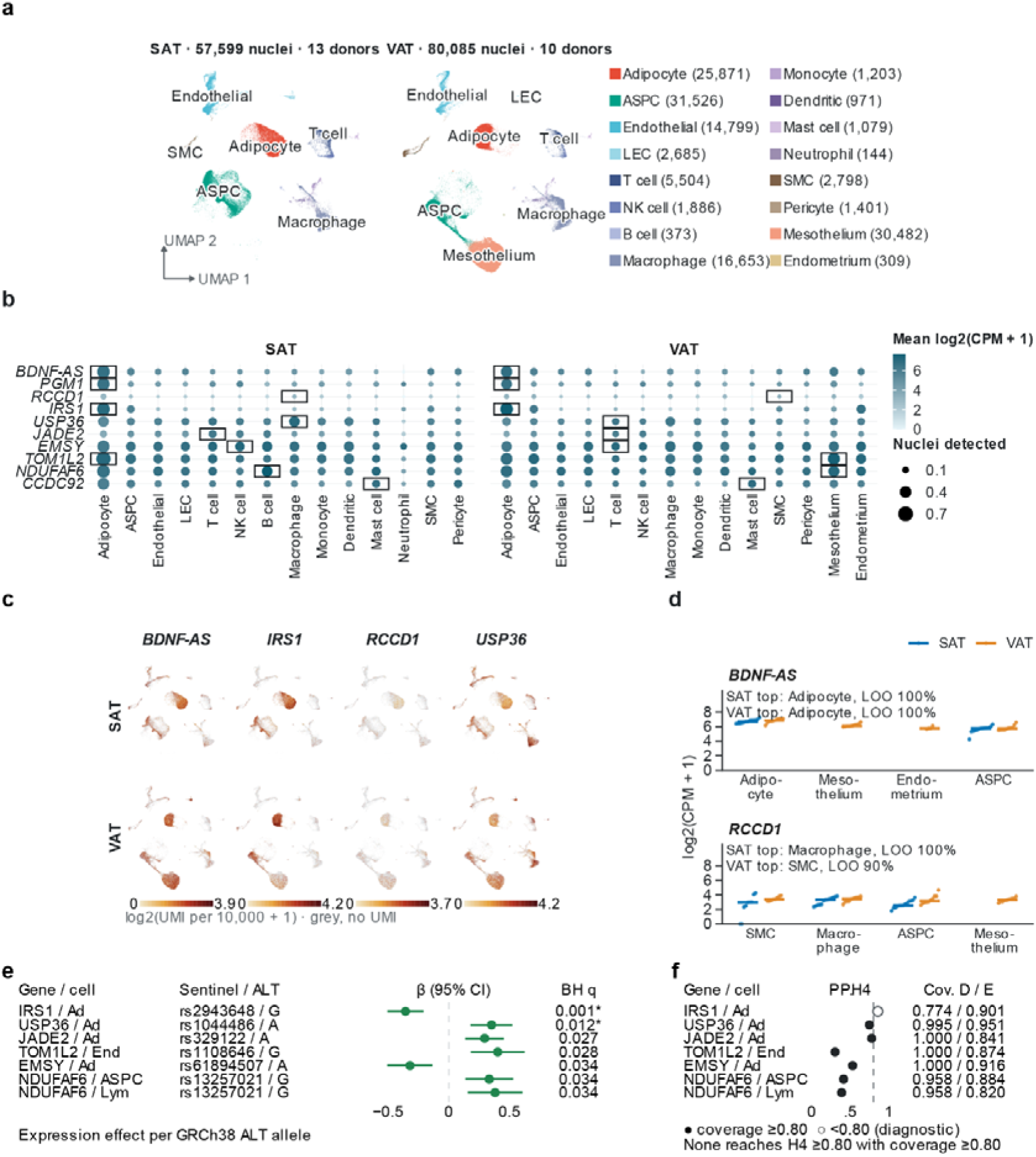
Cell-type context of prioritized adipose regulatory relationships. (a–d) Emont adipose snRNA-seq atlas (137,684 nuclei; 13 donors, ten also contributing VAT). (a) Author UMAP, one embedding shown by depot. (b) Genes resolved in Fig. 3, genes with associations in e, and CCDC92 (all 38 in Supplementary Fig. S7). Colour, equal-donor mean log2(CPM + 1); area, donor-averaged fraction of nuclei with detected expression; box, highest-mean cell type per depot; ×, <3 eligible donors. (c) Nucleus-level expression; colour scale shared by depot within a gene, capped at the 99th percentile of expressing nuclei. (d) Donor pseudobulks in each gene’s four highest-mean cell types; bars, medians; LOO, leave-one-donor-out retention of the top rank. Expression is gene-level, not the discovery RNA features. (e,f) KOBS/RYSA SAT cell-class eQTLs (separate cohort): (e) the seven BH-supported associations among 190 planned tests, GRCh38 ALT effects, unadjusted 95% intervals; (f) their regional models (BF*, prior SD 0.15, p12 = 5e−6); D/E, disease/expression coverage. Ad, adipocyte; End, endothelial; Lym, lymphoid.

For display, nuclei were placed on the authors’ published UMAP embedding retrieved from the MATKP portal; the coordinates covered exactly the 137,684 analysed nuclei with matching cell-type labels, and no re-clustering or new embedding was performed (Fig. 5a). Nucleus-level expression of display genes was shown as log□(UMI per 10,000 + 1) from the released count matrix (Fig. 5c), and the per-gene sums over nuclei reproduced every donor-level pseudobulk count.

### Cell-class expression QTLs and regional models

The SAT KOBS/RYSA cell-type eQTL release supplied nominal statistics for adipocytes, adipose stem and progenitor cells, endothelial cells, lymphoid cells and myeloid cells ^49^. Its source filters included a minor-allele frequency of at least 0.10 and a window extending 500 kb from gene ends. The original sentinel–gene combinations of the 38 relationships in the five cell classes defined 190 planned tests. Exact coordinate and allele matching aligned reported effects to the GRCh38 alternative allele. BH and Bonferroni corrections used the full planned denominator of 190, and unavailable statistics remained missing (Fig. 5e; Supplementary Fig. S8a; Table S8).

Regional models compared the disease signal with each cell-class expression profile in 30 branches: two disease representations (BF* or raw BF), three molecular-effect prior SDs (0.10, 0.15 and 0.20) and five joint priors. Molecular approximate BFs used *V* = SE^2^ and *W* = SD^2^, logBF = 1⁄2[log{*V*/*V* + *W*)} + *β*^2^*W*/ {*V*(*V* + *W*)}]^28^. The primary branch used BF*, SD 0.15 and *p*_12_ = 5 × 10^-6^. Eligibility required source integrity, at least two shared variants and bilateral coverage of at least 0.80; computable models failing these criteria were reported as diagnostic (Fig. 5f; Supplementary Figs. S8 and S9; Table S8).

### Metabolic-trait colocalization

The trait and tissue colocalization rules were specified before inspection of the trait results, with one revision: the use of trait coverage as a gate was changed after the first run showed that it classified traits without a regional signal as not testable. Decision rules for the extension analyses (BMI adjustment, sensitivity analyses, additional brain regions, islets, multi-signal fine-mapping and SMR/HEIDI) were written before their data were examined. One secondary lead-variant comparison was added after inspection and is reported as such.

We used European-ancestry summary statistics for fasting glucose, BMI-adjusted fasting insulin, BMI-adjusted 2-h glucose and HbA1c from MAGIC ^21^; BMI, waist-to-hip ratio and waist-to-hip ratio adjusted for BMI from GIANT–UK Biobank ^32^; triglycerides, HDL cholesterol, LDL cholesterol and total cholesterol from GLGC ^31^; HOMA-IR from MAGIC 2010 ^33^; and adiponectin from ADIPOGen ^34^. Variants within 2.6 Mb of the 30 supported signals were mapped from GRCh37 to GRCh38, oriented to the GRCh38 reference allele and restricted to biallelic single-nucleotide variants. Signed z-scores were derived from the reported P values and effect signs, and trait evidence was summarized as Wakefield approximate BFs on the per-SD scale with a prior SD of 0.15 ^28^. Missing MAGIC allele frequencies were taken from the GIANT–UK Biobank European data at the same site and alleles; rows still lacking a frequency were excluded.

Disease BF* (signal level) and the log BFs of each retained adipose component (component level) were compared with trait evidence over shared variants using the discovery priors and bilateral coverage of at least 0.80. Calls were made in order: not testable when fewer than two variants were shared or disease or molecular coverage was below 0.80; shared genetic evidence when trait coverage was at least 0.80 and PP.H4 ≥ 0.80; no detected association when no trait variant reached P < 5 × 10□□; not testable when the trait signal lay outside the shared variants; and otherwise association without colocalization (Supplementary Fig. S10a).

To allow several trait signals per region, each signal span was fine-mapped for the 11 traits with genome-wide data using susie_rss in susieR 0.12.35 (L = 10; 95% credible sets; purity ≥ 0.5) with UK Biobank European LD matrices restricted to variants with minor-allele frequency of at least 0.01 and the median trait sample size ^35,36^. Variants with a kriging log-likelihood ratio above 2 and |z| > 2 were removed. Non-converged fits were refitted once with the LD matrix regularized to 0.99R + 0.01I, and fits in which all ten components formed credible sets were treated as LD mismatch. Regions without a pure credible set were classified as not fine-mappable, and the single-effect result was retained. Each trait credible-set component was compared with each RNA component and with the disease BF* under the same model and coverage rule (Supplementary Fig. S10c). In the relationship-by-trait summary, each cell took the strongest state across the relationship’s retained components, in the order shared under both models, shared under the single-effect model only, shared under the multi-signal model only, association without colocalization, no detected association and not testable. Single-effect support in a region without a usable trait credible set was flagged separately (Fig. 6a; Table S9). Sensitivity analyses used trait-effect prior SDs of 0.10 and 0.20 and GLGC data excluding UK Biobank.

**Figure 6.**
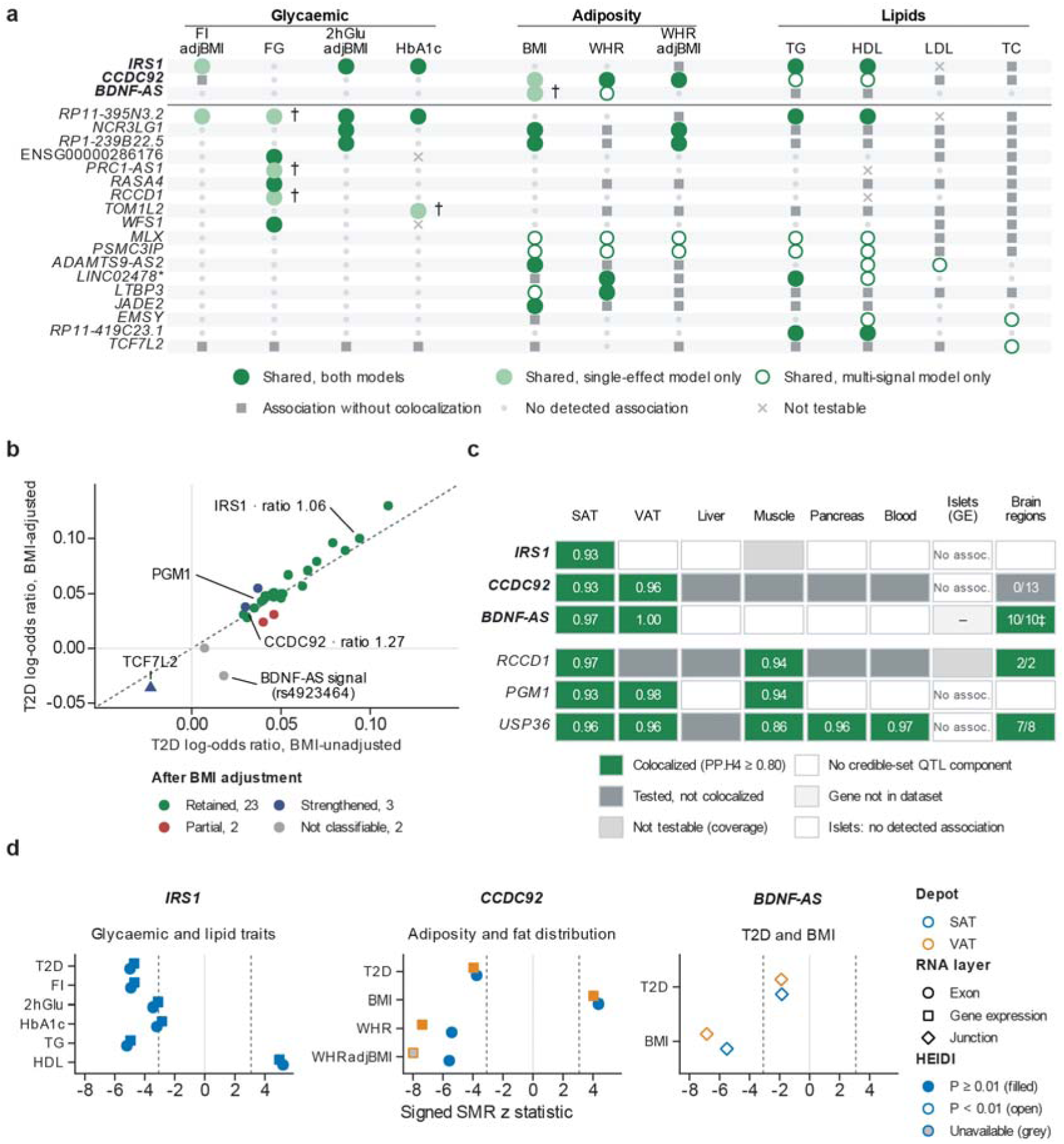
Metabolic-trait and tissue analyses define physiological contexts for prioritized adipose RNA relationships. (a) Trait profiles of the 21 relationships whose retained RNA components share genetic evidence (PP.H4 ≥0.80, coverage ≥0.80) with at least one of 13 traits (11 shown), under single-effect and multi-signal (SuSiE-RSS) trait models; cells take the strongest state across components. IRS1, CCDC92 and BDNF-AS are shown first. (b) Sentinel T2D log-odds ratios without and with BMI adjustment (30 signals); dashed, equality; ratio, adjusted/unadjusted log-odds ratio. Retained, ratio 0.8–1.25; strengthened, >1.25; partial, 0.5–0.8; not classifiable, unadjusted |z| < 3. TCF7L2 (triangle), conditional secondary signal. (c) Colocalization with same-gene QTLs in GTEx tissues, islets (gene-level statistics) and 13 brain regions (colocalized/testable). ‡ The same BDNF-AS junction as in adipose tissue. (d) Exploratory SMR at each feature’s top cis variant; dotted lines, Bonferroni threshold for 24 tests; SMR does not establish causality. Complete states for all 38 relationships are in Table S10.

### Tissue context

All credible-set components overlapping each signal were evaluated in GTEx liver, skeletal muscle, pancreas and whole blood (five RNA layers), in hypothalamus (five layers) and in 12 further GTEx brain regions (gene expression and LeafCutter junctions) from eQTL Catalogue release 7 ^19,20^. SAT and VAT were re-evaluated under the same per-tissue rule. Same-feature comparisons required identical feature identifiers, or identical intron coordinates and strand for LeafCutter junctions. Pancreatic islet gene-expression summary statistics from InsPIRE were analyzed with the single-effect model ^37^. A relationship was supported in a tissue when an admissible comparison for the same gene reached PP.H4 ≥ 0.80. Relationships without support in any non-adipose tissue were grouped by what could be tested: GTEx non-adipose tissues, brain regions only, or islet gene-level data only. Absence of a credible-set component was recorded as no detected QTL rather than absence of regulation (Fig. 6c; Supplementary Fig. S10b; Table S9).

### BMI adjustment and effect direction

Sentinel T2D effects were compared between the BMI-unadjusted and BMI-adjusted European analyses of DIAGRAM 2018, aligned to the DIAMANTE 2022 risk allele ^1^. The ratio of adjusted to unadjusted log-odds ratios defined four classes: retained (0.8–1.25), strengthened (> 1.25), partially attenuated (0.5–0.8) and attenuated (≤ 0.5). Signals with an unadjusted |z| < 3 were not classified (Fig. 6b).

Because adjustment for a heritable covariate can induce collider bias, strengthened effects were not interpreted as evidence of BMI-independent mechanisms ^38^. The T2D-raising allele at each lead variant was taken from the fixed-effects estimates of Suzuki et al. ^39^. RNA effects used eQTL Catalogue alternative-allele estimates, and only effect signs were compared across depots, tissues and RNA layers.

### SMR and HEIDI

For the *IRS1*, *CCDC92* and *BDNF-AS* features, summary-data-based Mendelian randomization (SMR) used the top variant in each feature’s complete cis region ^40^. The heterogeneity in dependent instruments (HEIDI) test used up to 20 variants with QTL P < 1.57 × 10□³ and 0.05 < r² < 0.9 with the top variant, UK Biobank LD and a Satterthwaite approximation. In total, 24 tests were performed, and a Bonferroni threshold for 24 tests was used for display; HEIDI P < 0.01 was taken to indicate heterogeneity (Fig. 6d; Table S9). Because SMR at a single cis locus is not independent of colocalization, these analyses were treated as exploratory.

### Integrated evidence and pharmacological annotation

Each relationship was assigned to a measurement class according to the molecular layers of its retained pairs: GE only, GE plus at least one non-GE layer, or non-GE layers without a retained GE pair. These classes define whether initial measurement should target total gene abundance, total RNA together with the implicated feature, or the implicated RNA feature itself (Fig. 7a; Table S10). For five representative relationships, the retained RNA objects, cellular contexts and proposed readouts were compiled from the preceding analyses (Fig. 7c; Table S10); these readouts are proposed experiments and were not performed.

**Figure 7.**
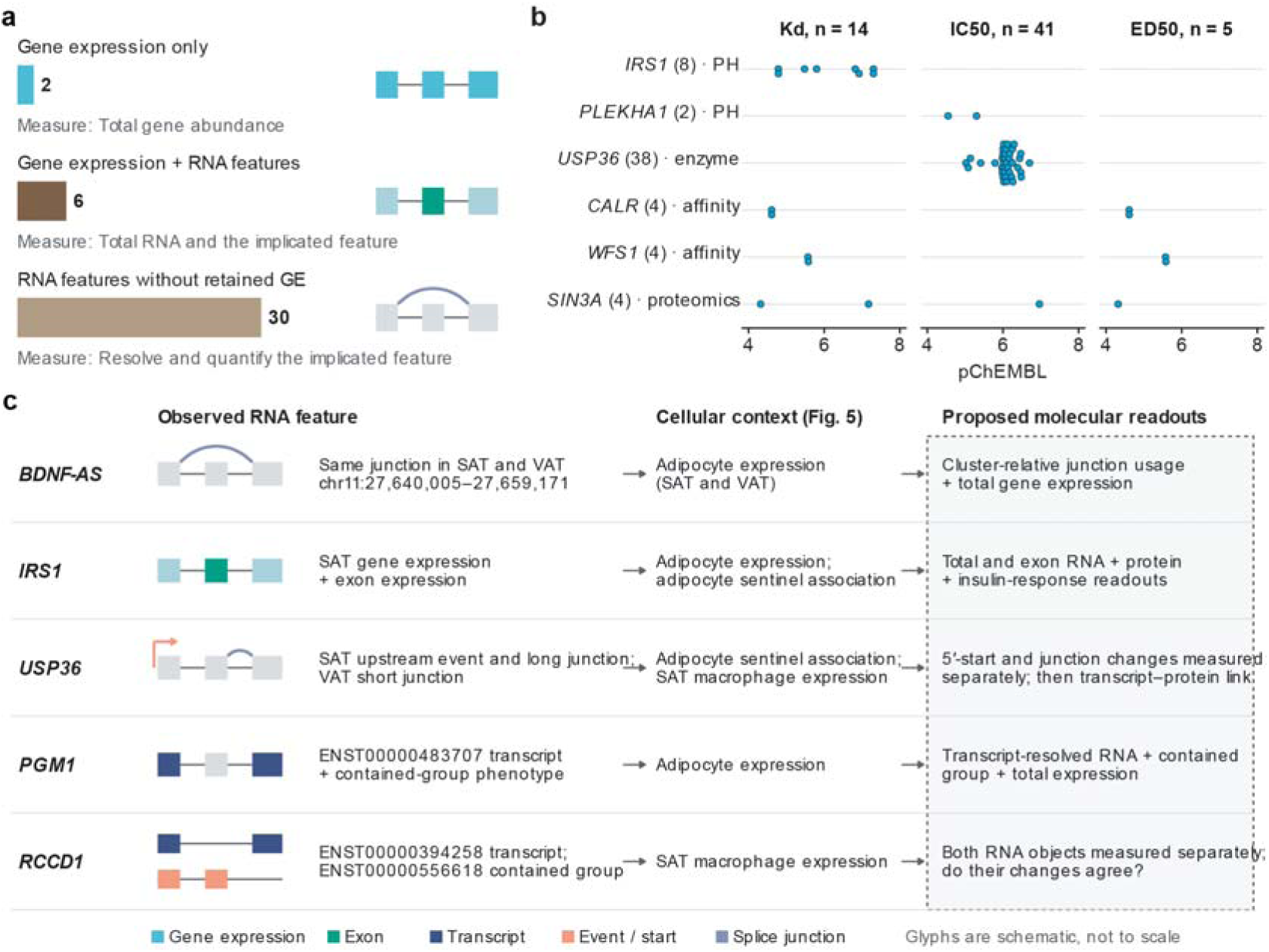
Integrated evidence specifies molecular measurements and experimental contexts for prioritized relationships. (a) Retained molecular evidence of the 38 relationships and the initial measurement it implies; categories follow the molecular layers in the retained atlas, and the 30 RNA-feature relationships span 25 signals. (b) Sixty quantitative ChEMBL activity records retained after target-identity review, shown by endpoint and experimental context; numbers in parentheses are records per relationship. Records are database-level assay observations, not independent validations, and are not ranked across endpoints. (c) Observed RNA features, cellular context from Figure 5 and proposed readouts (dashed box) for five representative relationships. Proposed readouts are follow-up experiments, not established mechanisms; arrows denote study design. Complete molecular objects, bases and readouts are in Table S10, and all activity records in Table S12.

All 38 relationships were annotated with Open Targets Platform 26.06, ChEMBL 37, the IUPHAR/BPS Guide to PHARMACOLOGY 2026.2 and DrugCentral release 54 ^41-44^. SwissProt-labelled protein accessions in Open Targets were checked against its release-matched tractability input. Positive tractability annotations were deduplicated by relationship within modality and evidence category. Essentiality and safety contexts were taken from DepMap common-essential lists, ToxCast and ClinPGx ^45-47^ and interpreted in their experimental context (Table S11).

All 126 ChEMBL activity records linked to the relationships were retained with assay, target, molecule, endpoint, units and data-quality fields. Quantitative records required an exact human single-protein target, a direct assay relationship, a binding or functional assay, a finite pChEMBL value and no data-validity comment. Review of the experimental object excluded eight records labelled MLX that measured an ML-IAP/XIAP chimera, leaving 60 records across six relationships. Kd, IC50 and ED50 records were kept separate and not compared across endpoints (Fig. 7b; Table S12). Because aligned signal-specific disease and component-consistent molecular effects were not available for all relationships, proposed follow-up specified molecular readouts and cellular contexts without prescribing activation or inhibition.

### Statistical analysis

Colocalization evidence was summarized as posterior probabilities; the discovery set was defined by the cumulative mean-score rule described above, and component-level comparisons used PP.H4 ≥ 0.80 together with bilateral coverage of at least 0.80. Multiple testing was controlled within each prespecified family using its full planned denominator: Bonferroni correction over eight METSIM fixed-variant tests, BH correction over six KOBS lookups, BH and Bonferroni corrections over 190 cell-class tests, and a Bonferroni display threshold over 24 SMR tests. Unavailable statistics remained missing and were never treated as null results. Effect estimates are reported with unadjusted 95% confidence intervals where shown. No hypothesis tests were performed for descriptive summaries of testability, expression or annotation.

### Software and reproducibility

The primary analyses used R 4.5.2 with coloc 6.0.1, together with Python 3.11 (NumPy, pandas and SciPy) and R 4.4.2 with susieR 0.12.35 for multi-signal fine-mapping. An independent re-implementation of the colocalization model reproduced all 77 retained pairs with identical shared sets and a maximum PP.H4 difference of 3 × 10□¹□. OpenAI ChatGPT assisted with language editing.

## Results

### Multi-layer RNA integration expands T2D signal testability and defines an adipose regulatory atlas

To map type 2 diabetes (T2D) genetic signals to adipose RNA phenotypes, we integrated 338 fine-mapped T2D signals from DIAMANTE with GTEx molecular quantitative trait loci (QTLs) uniformly processed by the eQTL Catalogue^2,19,20^. The analysis encompassed gene expression (GE), exon expression, transcript usage, transcriptional-event usage and splice-junction usage^16,48^ in subcutaneous (SAT) and visceral adipose tissue (VAT), providing ten depot–layer datasets within a common analytical framework (Fig. 1a).

Multi-layer integration expanded the range of T2D signals available for adipose colocalization analysis. Of 50,995 candidate signal–component pairs, 13,216 met the shared-variant posterior coverage requirement of at least 0.80 on both the disease and molecular sides, together with component and variant integrity checks (Fig. 1b; Supplementary Fig. S1). A signal was considered testable when it had at least one admissible comparison. Integrating all five layers increased the number of testable signals from 253 with GE alone (74.9%) to 293 (86.7%), adding 40 signals (Fig. 1c; Supplementary Fig. S2a,b; Table S2). This gain extended across multiple RNA measurements: 28 of the 40 additional signals (70.0%) were testable in at least two non-GE layers (Supplementary Fig. S2c).

Applying a cumulative mean-score selection rule across all 13,216 admissible comparisons selected 77 pairs, with a mean posterior non-colocalization score of 0.04985, below the selection limit of 0.05 (Fig. 1d). All 77 pairs were retained after signal attribution, with no competition group containing more than one selected disease signal (Supplementary Fig. S3d). Grouping the retained pairs by signal and gene defined 38 signal–gene relationships across 30 T2D signals (Table S1). Across the complete set of 338 input signals, 30 had at least one retained colocalization pair, 263 were tested without detected support and 45 were not testable (Fig. 1e).

Both adipose depots contributed to the atlas. Colocalization support was detected for 23 signals in SAT and 21 in VAT, comprising 14 supported in both depots, nine in SAT only and seven in VAT only (Fig. 1f). These single-depot detections occurred at signals that were testable in both depots. The relationship-level atlas recorded the RNA layers and molecular components contributing to each signal– gene relationship across SAT and VAT (Fig. 1g).

The primary pair set was nested between the sets obtained with fixed PP.H4 thresholds. PP.H4 > 0.90 selected 71 pairs representing 35 relationships, all contained within the primary set, whereas PP.H4 > 0.80 selected 110 pairs representing 59 relationships and included all 77 primary pairs (Supplementary Fig. S3b,c). Three component-thinning schemes, each retaining one representative component per resolvable molecular phenotype and repeating selection and attribution, preserved 33–36 of the 38 relationships without identifying additional relationships (Supplementary Fig. S3e). All 77 pairs were also retained in each of five perturbations within the rounding intervals of the released disease posteriors (Table S3).

Together, multi-layer integration expanded the testable T2D signal space and established an adipose RNA atlas of 38 signal–gene relationships, providing a basis for examining the molecular information contributed beyond total gene expression.

### Non-gene-expression measurements identify specific RNA features at T2D signals

To examine the molecular information contributed beyond total gene expression, we assessed retained colocalization evidence across RNA layers. Of the 77 retained component pairs, 67 (87.0%) involved non-gene-expression (non-GE) measurements: exon expression (20 pairs), transcript usage (15), transcriptional-event usage (19) and splice-junction usage (13), compared with ten GE pairs (Fig. 2a). Non-GE measurements also accounted for 86.0% of admissible comparisons, with similar observed retention proportions for GE and non-GE comparisons (0.54% and 0.59%, respectively). Across the 30 supported signals, the layers contributed overlapping sets of detections, with individual signals supported through different combinations of RNA measurements and adipose depots (Fig. 2b).

We next compared non-GE and GE evidence at signals where total expression could also be evaluated. Of the 30 supported signals, 23 (76.7%) had at least one retained non-GE pair but no retained GE pair. Within this subset, nine signals had an admissible but unselected GE comparison for at least one implicated gene in either depot; four had GE comparisons that did not meet the coverage requirement, and ten lacked a comparable GE feature (Fig. 2c). The nine signals with evaluable GE evidence formed the subset for detailed comparison.

At these nine signals, 16 retained non-GE component pairs represented 11 signal–gene relationships and 13 distinct molecular features. Admissible same-gene GE comparisons were available for 14 pairs spanning nine relationships; NCR3LG1 lacked a same-gene GE feature, whereas the PRC1-AS1 GE comparison did not meet the coverage requirement. For each of the 14 comparable pairs, we selected the admissible same-signal, same-gene GE comparison with the highest PP.H4 across either depot as the representative, recording its depot to distinguish same-depot from cross-depot contrasts (Fig. 2d; Supplementary Table S4).

At BDNF-AS and PGM1, specific RNA features colocalized with T2D while the evaluated GE component favored distinct disease and expression variants. The same BDNF-AS splice junction colocalized in SAT and VAT (PP.H4 = 0.972 and 0.997, respectively), whereas the VAT GE component favored distinct variants (PP.H3 = 0.999). At PGM1, retained transcript-usage and transcriptional-event components yielded PP.H4 values of 0.895–0.977, while the SAT GE component likewise favored distinct variants (PP.H3 = 0.9996; Fig. 2d).

At SAXO1 and RCCD1, GE also showed shared-variant support alongside retained evidence for specific RNA features. At SAXO1, SAT event usage and SAT GE had PP.H4 values of 0.890 and 0.837, respectively, although only event usage was retained under the primary selection rule. At RCCD1, SAT transcript-usage and transcriptional-event components reached PP.H4 = 0.957 and 0.969, compared with 0.782 for the VAT GE representative (Fig. 2d). The admissible same-depot SAT GE component L2 instead favored distinct variants (PP.H3 = 0.9969; Supplementary Fig. S4m,n).

To guide follow-up across the full atlas, we summarized retention of all 38 relationships under lower joint priors and, in separate reruns, higher bilateral coverage requirements (Fig. 2e; Supplementary Table S4). Four relationships—BDNF-AS, JADE2, ANK1 and TCF7L2—were retained at the lowest tested joint prior (p12 = 5 × 10⁻□). Among these, BDNF-AS and JADE2 were also retained in the separate analysis requiring bilateral coverage of at least 0.95.

Within the 11 relationships represented in Fig. 2d, seven were retained at a bilateral coverage requirement of 0.90 and three—BDNF-AS, PGM1 and RCCD1—at 0.95. All three were also retained at p12 = 2 × 10⁻□ and with raw disease Bayes factors. BDNF-AS was the only member of this subset retained across all evaluated sensitivity branches (Supplementary Table S4). These three relationships were therefore selected for locus-resolved analysis of their RNA features and conditional candidate variants.

### Locus-resolved colocalization links RNA features to candidate variants at BDNF-AS, PGM1 and RCCD1

We resolved the three most robust relationships—BDNF-AS, PGM1, and RCCD1—to specific RNA structures and conditional candidate variants. At the rs4923464 T2D signal, the retained SAT and VAT splicing components mapped to the identical BDNF-AS junction (chr11:27,640,005–27,659,171, GRCh38) (Fig. 3a). Junction usage colocalized with T2D in both depots, with PP.H4 = 0.972 in SAT and 0.997 in VAT (Fig. 3b–d).

Conditional on the shared-variant hypothesis (H4), rs6265 emerged as the leading candidate in both depots, with posterior probabilities of 0.813 in SAT and 0.997 in VAT (Fig. 3e). The SAT 95% conditional candidate set contained ten variants, whereas the VAT set contained rs6265 alone, making it the only variant shared by both depots (Supplementary Table S5). Notably, while rs4923464 was the reported sentinel, rs6265 actually led the T2D fine-mapping posterior (probability = 0.494; Fig. 3b).

The narrower VAT candidate set reflected the more concentrated molecular posterior at rs6265 in VAT compared to SAT (0.849 versus 0.097; Fig. 3c,d). In contrast, the evaluated VAT gene-expression component peaked 70–80 kb away (27.58–27.59 Mb) and strongly favored distinct variants (PP.H3 = 0.9993; PP.H4 = 3.1 × 10⁻□; Fig. 3f), clearly distinguishing the disease-associated junction signal from total BDNF-AS expression.

At the PGM1 signal (rs11576729), three retained measurements—SAT transcript usage, SAT transcriptional-event usage, and VAT transcriptional-event usage—shared a common transcript anchor (ENST00000483707). Their PP.H4 values were 0.933, 0.895, and 0.977, respectively (Supplementary Fig. S4a–d). The corresponding 95% conditional candidate sets contained ten, eleven, and seven variants, with five variants shared across all three (Supplementary Fig. S4e; Supplementary Table S5). The VAT event comparison was led by rs2269239 (which also led the T2D fine-mapping posterior at 0.327), whereas both SAT comparisons were led by rs3790858, which lay outside the VAT set. These measurements thus localized to overlapping candidate sets around a common transcript anchor, with leading variants diverging across depots.

At the RCCD1 signal (rs2890156), measurements with different transcript anchors produced nested candidate sets. SAT transcript usage anchored to ENST00000394258 (PP.H4 = 0.957), while SAT transcriptional-event usage anchored to ENST00000556618 (PP.H4 = 0.969; Supplementary Fig. S4h– k). All five variants in the event set were nested within the seven-variant transcript set, and both sets included the sentinel rs2890156, which also led the T2D fine-mapping posterior (0.334). The transcript and event comparisons were led by rs8028409 and rs79548680, respectively, and both leading variants belonged to both sets (Supplementary Fig. S4l; Supplementary Table S5). The two RNA measurements therefore prioritized the same candidate group while assigning different posterior weights within it.

These locus-resolved analyses reveal three distinct modes of molecular resolution: a single leading variant shared across depots for a BDNF-AS junction, overlapping candidate sets for PGM1 measurements anchored to a common transcript, and nested candidate sets for RCCD1 measurements with different anchors. These modes provide concrete, testable hypotheses for targeted functional experiments.

### External adipose datasets validate prioritized relationships and distinguish splicing from expression signals

To assess external support, we evaluated the prioritized relationships using adipose molecular data from FUSION, TwinsUK, METSIM, and KOBS^18,26,27,29^. FUSION and TwinsUK provided component-level colocalization data, METSIM provided fixed-variant junction associations and regional splicing/expression profiles, and KOBS provided sentinel–gene expression statistics (Fig. 4; Supplementary Table S6).

In FUSION and TwinsUK, 12 of the 46 evaluated signal–feature targets had an admissible component-level comparison in at least one cohort. Ten received retained colocalization support, representing eight signal–gene relationships (Fig. 4a), with LINC02478* and MLX each supported through two molecular features. In METSIM, statistics were available for five of the eight planned fixed-variant junction tests. Three associations passed Bonferroni correction: the USP36 junction (adjusted P = 8.57 × 10□²□), the LINC02478*-mapped junction (Padj = 1.23 × 10□□), and the longer PCGF3 junction (Padj = 1.34 × 10□□; Fig. 4b). Target-specific statistics were unavailable for BDNF-AS, MPHOSPH9, and ADAMTS9-AS2.

Two contrasting patterns of regional support emerged at USP36 and WFS1 (Fig. 4c). At USP36, junction usage strongly supported a shared variant with T2D (PP.H4 = 0.953), whereas total expression favored distinct variants (PP.H3 = 0.805; PP.H4 = 0.195). Conversely, at WFS1, gene expression supported a shared variant (PP.H4 = 0.944), while the examined junction lacked a clear regional QTL signal (PP.H1 = 0.768). Both contrasts persisted when restricting to the common variant set. Beyond these loci, the LINC02478*-mapped junction also showed regional colocalization with T2D (PP.H4 = 0.949). At PCGF3, however, the significant fixed-variant association coexisted with distinct-variant regional profiles for both junction and gene expression (PP.H3 > 0.999).

These regional signals proved highly robust (Supplementary Fig. S5). The USP36 and LINC02478* junction profiles, along with WFS1 gene expression, retained PP.H4 ≥ 0.80 at lower joint priors (p12 ≥ 2 × 10□□), under alternative disease Bayes factors, molecular-effect priors, and minor-allele-frequency filtering. Support diminished only at the most restrictive priors, and palindrome exclusion reduced QTL-side coverage below 0.80 for USP36 splicing and WFS1 expression.

Finally, KOBS provided sentinel–gene expression statistics for four of six planned tests, all passing Benjamini–Hochberg correction (PCGF3, RP11-395N3.2, IRS1, and ANK1; Fig. 4d; Table S6). Notably, PCGF3 received gene-expression support in FUSION/TwinsUK and KOBS, yet its METSIM regional profiles favored distinct variants, illustrating how evidence varies by resolution.

Together, these independent cohorts validate the prioritized relationships and establish that the informative molecular measurement—whether splice-junction usage or total expression—diverges substantially by locus (as seen at USP36 versus WFS1). This reinforces the fundamental necessity of multi-layer profiling to capture the specific genetic signals underlying T2D.

### Single-nucleus expression and cell-type genetic associations define cellular contexts for prioritized relationships

To identify cellular settings for follow-up, we examined the 38 prioritized relationships in the Emont human adipose single-nucleus atlas^9^. The analysis included 137,684 nuclei from 13 donors, with SAT available from all 13 and VAT from ten of the same donors (Fig. 5a). Thirty-five candidate genes mapped to the expression resource. Expression was quantified as log□(CPM + 1) within donor–depot– cell-type pseudobulks and averaged across eligible donors with equal weights. Within-donor nuclear detection fractions were summarized in parallel (Fig. 5b; Supplementary Fig. S7; Supplementary Table S7).

Across the 70 gene–depot combinations, adipocytes were the most frequent highest-expression cell type (13), followed by T cells (10) and NK cells (8). Eighteen genes had the same highest-expression cell type in both depots, and 13 retained their top rank in every leave-one-donor-out analysis in both depots (Supplementary Fig. S7b; Supplementary Table S7). BDNF-AS, PGM1 and IRS1 had their highest donor-averaged expression in adipocytes in both depots, with stable rankings after donor deletion.

RCCD1 showed a different distribution, with the highest mean expression in SAT macrophages and VAT smooth muscle cells (90% donor-deletion retention in VAT; Fig. 5b,d; Supplementary Fig. S7a; Supplementary Table S7). These profiles placed the locus-resolved candidates in distinct adipocyte, immune, and vascular expression backgrounds.

We next assessed the original T2D sentinel–gene combinations across five SAT cell classes using KOBS/RYSA eQTL statistics^49^. Of 190 planned sentinel–gene–cell-class tests, 68 had available statistics, and seven associations across six genes passed Benjamini–Hochberg correction over the full planned family (Fig. 5e; Supplementary Table S8). Four associations occurred in adipocytes, involving IRS1, USP36, JADE2 and EMSY. TOM1L2 was associated in endothelial cells, and NDUFAF6 in both adipose stem/progenitor cells and lymphoid cells. The IRS1 and USP36 adipocyte associations also passed Bonferroni correction across all 190 tests (adjusted P = 0.0013 and 0.023, respectively).

Regional models provided the highest shared-variant posterior for adipocyte IRS1 (PP.H4 = 0.857), followed by JADE2 (0.777) and USP36 (0.748). Disease-side coverage for IRS1 was 0.774 (diagnostic only), whereas the other six associations met the bilateral coverage requirement but had PP.H4 values of 0.303–0.777. None of the seven associations met both criteria for colocalization (Fig. 5f; Supplementary Fig. S8; Supplementary Fig. S9; Supplementary Table S8). These models therefore demarcate the limits of gene-level sentinel associations in specific cell types, highlighting that while the cellular measurement context is established, the specific RNA features must be directly quantified within these backgrounds to capture the disease-aligned regulatory signals.

Together, the integration of single-nucleus expression and cell-type genetics defined adipocytes (for BDNF-AS, PGM1, IRS1, and USP36) and macrophages/smooth muscle cells (for RCCD1) as primary cellular contexts. Notably, IRS1 combined a dominant adipocyte expression background with an adipocyte-specific genetic association, whereas USP36 showed an adipocyte genetic association but higher average expression in SAT macrophages, identifying complementary cellular settings for subsequent molecular investigation.

### Metabolic-trait and tissue analyses distinguish distinct physiological profiles of prioritized adipose RNA relationships

To link the prioritized adipose RNA features to metabolic phenotypes, we compared the 77 retained components with 13 European-ancestry traits, using single-effect models complemented by multi-signal analyses of 11 traits^21,31-34^. Shared genetic evidence was identified for 105 component–trait pairs spanning 21 of the 38 signal–gene relationships under at least one model (Fig. 6a; Supplementary Fig. S10a,c; Supplementary Table S9). We combined these trait profiles with BMI-adjusted T2D effects and cross-tissue colocalization; IRS1, CCDC92, and BDNF-AS illustrate three contrasting physiological contexts.

The IRS1 relationship showed an insulin-resistance and dyslipidaemia profile. Its SAT gene-expression and exon-expression components colocalized with BMI-adjusted 2-h glucose (PP.H4 = 0.92–0.93), HbA1c (0.85–0.87), triglycerides (0.96), and HDL cholesterol (0.94). The T2D risk allele was associated with lower adipose IRS1 expression and a worse metabolic profile. Importantly, its T2D effect estimate was essentially unchanged by BMI adjustment^1^ (ratio 1.06; Fig. 6b), and exploratory SMR associated the RNA features with T2D, fasting insulin, triglycerides, and HDL cholesterol, without detected HEIDI heterogeneity (P_HEIDI ≥ 0.12; Fig. 6d). IRS1 components were not testable in non-adipose GTEx or brain datasets, and islet gene-expression data showed no detected association (Fig. 6c) ^37^.

The CCDC92 relationship showed a fat-distribution profile. Its SAT exon and VAT gene-expression components shared genetic evidence with waist-to-hip ratio adjusted for BMI (WHRadjBMI; PP.H4 = 0.997 and 0.996). The T2D risk allele was associated with lower CCDC92 expression, higher waist-to-hip ratio, and lower BMI. In stark contrast to IRS1, BMI adjustment strengthened the CCDC92 T2D effect (ratio 1.27; Fig. 6b), a pattern consistent with limited peripheral fat storage ^38,50,51^. This relationship was highly tissue-restricted: colocalization was detected in both adipose depots, but not in liver, skeletal muscle, pancreas, whole blood, or any of 13 brain regions, despite testable components in all of them (Fig. 6c).

The BDNF-AS junction combined an adiposity-related trait profile with cross-tissue sharing. Junction usage colocalized with BMI in SAT and VAT (PP.H4 = 0.98 and 1.00) and with waist-to-hip ratio in the multi-signal analysis, but not with glycaemic or lipid traits. Unlike IRS1 and CCDC92, this junction was not adipose-restricted: it colocalized with the T2D signal in all ten testable GTEx brain regions, including the hypothalamus (PP.H4 = 0.92–1.00; Fig. 6c; Supplementary Fig. S10b). While SMR associated junction usage with BMI in both depots, HEIDI detected heterogeneity (P_HEIDI = 1.6 × 10□□ in SAT and 3.0 × 10□¹² in VAT; Fig. 6d), suggesting that the cross-tissue genetic signal at this locus is complex and may involve multiple linked regulatory variants^40^.

Other locus-resolved relationships showed distinct trait patterns. RCCD1 transcript and event components shared evidence with fasting glucose (PP.H4 = 0.90 and 0.92), although the regional trait association did not reach genome-wide significance. Neither PGM1 nor USP36 shared genetic evidence with an examined testable trait (Table S9). Across the full atlas, 21 relationships colocalized with same-gene QTLs in at least one non-adipose tissue, eight of which also had metabolic-trait support. In all 70 cross-depot or cross-tissue comparisons in which the same RNA feature colocalized, molecular-effect directions were strictly concordant when aligned to the T2D risk allele. Trait-effect prior sensitivity analyses preserved all 28 signal-level and 51 of 52 component-level single-effect colocalization calls, and 12 of 13 component-level lipid calls were recovered using GLGC data excluding UK Biobank (Table S9).

### Integrated evidence specifies molecular measurements and cellular settings for follow-up

To translate the 38 prioritized relationships into testable experiments, we integrated their molecular features, external support, and cellular profiles with existing pharmacological annotations. Based on the layers represented in the retained atlas, these relationships fell into three measurement classes: two supported by gene expression alone, six combining gene-expression and RNA-feature support, and 30 supported by RNA features without a retained gene-expression pair (Fig. 7a; Table S10). This classification dictates the initial experimental readout: total gene abundance, paired measurement of total and RNA-feature expression, or direct resolution of the implicated RNA feature.

Pharmacological and tractability annotations provided the experimental context for these targets (Fig. 7b; Tables S11 and S12) ^41-44^. SwissProt-labelled protein accessions were available for 27 relationships, and 60 quantitative ChEMBL activity records were retrieved across six relationships, including enzyme assays for USP36 (38 records) and PH-domain binding assays for IRS1 (eight records). Additional evidence from DepMap, ToxCast, and ClinPGx highlighted essentiality and pharmacogenomic contexts for specific targets (e.g., USP36 and SIN3A in the DepMap common-essential list) ^45-47^. These annotations define the biochemical starting points for designing targeted assays.

Combining these annotations with the identified RNA features and cellular profiles specified distinct follow-up measurements for five representative relationships (Fig. 7c). For BDNF-AS, the proposed readout pairs cluster-relative usage of the chr11:27,640,005–27,659,171 junction with total BDNF-AS expression in SAT and VAT adipocyte backgrounds. For IRS1, its combined gene-expression and exon evidence, adipocyte sentinel association, and metabolic profile motivate paired measurement of total and exon RNA alongside IRS1 protein abundance and insulin-response readouts in adipocytes. For USP36, the distinct SAT upstream transcriptional event, longer SAT junction, and shorter VAT junction require separating 5’-start and junction changes in SAT adipocytes before resolving their protein consequences. For PGM1, the priority is transcript-resolved quantification of ENST00000483707 and its contained-group phenotype alongside total expression. For RCCD1, transcript usage of ENST00000394258 and the ENST00000556618 contained group define two separate RNA objects for independent measurement in SAT macrophages.

Together, these routes connect the genetic observations to measurable molecular consequences. The resulting atlas provides specific variants, RNA features, and cellular backgrounds for direct experimental interrogation of how adipose regulation contributes to T2D susceptibility.

## Discussion

Integrating five complementary RNA measurements across subcutaneous and visceral adipose tissue, we constructed a multi-layer regulatory atlas linking 30 fine-mapped T2D signals to 38 signal–gene relationships. Rather than only expanding the catalog of nominated candidate genes, this framework resolves the specific molecular features—exon expression, transcript usage, transcriptional-event usage, and splice-junction usage—through which T2D genetic signals can be resolved in adipose tissue. By uniting component-level colocalization with four external adipose cohorts, single-nucleus transcriptomics, metabolic-trait genetics, and pharmacological annotations, our study distinguishes loci supported by overall transcript abundance from those requiring isoform- or processing-level measurements. This resolution sharpens both the biological interpretation of T2D susceptibility loci and the experimental readouts required for functional validation.

Previous studies have established that genetic effects on transcript usage and splice-junction regulation contribute substantially to complex traits and cardiometabolic phenotypes, often operating independently of total gene expression^15,17,18,20^. Uniformly processed resources such as the eQTL Catalogue have enabled systematic evaluation of these regulatory layers ^19^. Building on these foundations, our signal-level integration across two adipose depots demonstrates the practical power of multi-layer profiling. Incorporating four transcript-processing layers alongside gene expression expanded the number of testable T2D signals from 253 to 293 (86.7% of all fine-mapped signals), and the four non-GE layers contributed 86.0% of admissible comparisons, expanding the comparison pool more than sixfold relative to gene expression alone. At the same time, 23 of the 30 supported T2D signals (76.7%) had retained non-GE evidence without a retained GE pair. The added value of multi-layer analysis is clearest at loci with direct same-gene comparators, such as BDNF-AS and PGM1, where specific splice-junction or transcript features strongly colocalized with T2D (PP.H4 = 0.895– 0.997) while total gene expression at the same locus favored distinct causal variants (PP.H3 > 0.999). These direct contrasts show that relying on bulk gene-level expression alone can miss or mischaracterize the regulatory signal captured by feature-level measurements at established disease loci.

Locus-resolved fine-mapping of BDNF-AS, PGM1, and RCCD1 illustrates three distinct modes by which feature-level analysis sharpens mechanistic hypotheses. At the BDNF/BDNF-AS locus, usage of a single BDNF-AS splice junction (chr11:27,640,005–27,659,171) colocalized with T2D in both SAT and VAT, converging on rs6265 as the leading candidate variant conditional on H4 (81.3% posterior mass in SAT and 99.7% in VAT), whereas VAT BDNF-AS gene expression peaked 70–80 kb away and supported distinct variants. This finding bridges two previously separate lines of biological evidence: rs6265 encodes the well-characterized BDNF Val66Met substitution that alters activity-dependent BDNF secretion^52^, while antisense transcription of BDNF-AS is known to repress endogenous BDNF expression^53^. Pinpointing a specific BDNF-AS splicing event aligned with rs6265 suggests that coding variation and antisense RNA processing may be coupled at this locus, supporting functional perturbations that target the specific splice junction alongside total BDNF-AS abundance.

Complementing this single-variant convergence, PGM1 and RCCD1 demonstrate how multiple transcriptomic layers narrow candidate variant sets in more complex LD architectures. At PGM1, SAT transcript usage and SAT/VAT transcriptional-event measurements converged on a shared transcript anchor (ENST00000483707) and five core candidate variants across depots; at RCCD1, measurements anchored to distinct transcripts (ENST00000394258 and ENST00000556618) produced nested five- and seven-variant candidate sets containing the T2D sentinel rs2890156. Together, these patterns narrow broad association intervals to specific RNA objects and compact candidate-variant sets for targeted perturbation.

Evaluation across four independent adipose cohorts—FUSION, TwinsUK, METSIM, and KOBS— provided complementary support for prioritized relationships and showed that the informative molecular measurement can differ by locus ^18,26,27,29^. In FUSION and TwinsUK, ten of the 12 targets with an admissible component-level comparison retained colocalization support across eight signal– gene relationships, including dual-feature support for LINC02478* and MLX. In METSIM, regional profiling showed a sharp functional dichotomy: the USP36 splice junction strongly colocalized with T2D (PP.H4 = 0.953) while its total gene expression favored distinct variants (PP.H3 = 0.805), whereas WFS1 colocalized robustly through gene expression (PP.H4 = 0.944) rather than splicing. At the same time, the coexistence of a significant fixed-variant association with distinct regional variants (PP.H3 > 0.999) at PCGF3 in METSIM illustrates the analytical advantage of full regional colocalization over single-variant lookups in distinguishing shared signals from LD hitchhiking.

Integrating the Emont single-nucleus atlas with KOBS/RYSA cell-class eQTLs placed these molecular relationships into defined cellular environments^9,49^. Single-nucleus expression profiles identified mature adipocytes as the predominant expression setting for BDNF-AS, PGM1, and IRS1 in both depots, while pointing to SAT macrophages and VAT smooth muscle cells for RCCD1. In parallel, cell-class eQTL mapping identified seven BH-significant sentinel associations across six genes, including adipocyte associations for IRS1, USP36, JADE2, and EMSY, an endothelial association for TOM1L2, and progenitor/lymphoid associations for NDUFAF6. Notably, IRS1 combined dominant adipocyte expression with the highest regional H4 in adipocytes (PP.H4 = 0.857) in a diagnostic model, nominating the adipocyte context for feature-level follow-up, whereas USP36 showed an adipocyte-specific genetic association despite higher baseline expression in macrophages. Moreover, the moderate posterior probabilities from gene-level regional models in sorted cell classes (PP.H4 = 0.303–0.777 among coverage-eligible tests) reinforce a central finding of our bulk analysis: because the prioritized signals often involve specific exons, junctions, or transcript events, capturing their regulatory signal at the cellular level will require single-cell or nucleus-sorted isoform and splicing quantification.

Linking the adipose RNA atlas to 13 metabolic traits, BMI-adjusted T2D summary statistics, and multi-tissue QTLs delineated three distinct pathophysiological contexts among the prioritized loci. First, IRS1 exemplified an adipocyte insulin-resistance and dyslipidaemia profile: T2D risk alleles associated with lower adipose IRS1 expression, adverse glycaemic and lipid traits (2-h glucose, HbA1c, triglycerides, and HDL cholesterol), a BMI-invariant T2D effect (ratio 1.06), and consistent SMR support without HEIDI heterogeneity. This aligns with classic physiological evidence connecting reduced adipose IRS1 expression to impaired insulin signaling and dyslipidaemia^54-56^. Second, CCDC92 defined an adipose-restricted, fat-distribution context: its SAT exon and VAT expression components colocalized with WHRadjBMI, exhibited an amplified T2D effect after BMI adjustment (ratio 1.27), and showed an adipose-restricted colocalization pattern across the surveyed testable GTEx tissues and 13 brain regions. Together with functional studies implicating CCDC92 in adipocyte differentiation and lipid storage^50,57^, this profile points to a context involving peripheral subcutaneous fat expandability in T2D risk^50,51^. Third, the BDNF-AS splice junction represented an adiposity-related context with cross-tissue sharing, colocalizing with BMI and WHR and sharing the identical T2D-colocalized junction across SAT, VAT, and all ten testable GTEx brain regions, including the hypothalamus^58,59^. The presence of HEIDI heterogeneity at BDNF-AS suggests a complex local regulatory architecture potentially involving multiple linked variants across neural and adipose tissues.

A key translational contribution of this work is the conversion of statistical colocalizations into a practical, measurement-led experimental framework. Classifying the 38 relationships by their retained molecular layers reveals that 30 relationships cannot be evaluated by total expression alone; they require direct quantification of specific junctions, exons, or transcript events alongside total-expression controls. Combining this classification with cellular contexts and 60 curated ChEMBL bioactivity records—such as enzymatic assays for the deubiquitinase USP36 ^60^ and PH-domain binding assays for IRS1—provides concrete starting points for functional interrogation. For instance, follow-up of USP36 in adipocytes can pair separated 5’-start and splice-junction readouts with established deubiquitinase activity assays, whereas PGM1 and RCCD1 call for isoform-resolved quantification in adipocytes and macrophages, respectively, before protein-level characterization.

Several methodological considerations frame the scope of our findings and point to future extensions. First, our discovery atlas operates within the boundaries of currently released fine-mapping credible sets and bilateral posterior coverage (≥0.80); loci failing coverage gates or lacking annotated features represent unassayed space that may be resolved with denser reference panels and long-read transcriptomic catalogs. Second, cross-cohort differences in sample size, isoform annotation, and junction quantification denominators mean that external datasets provide complementary support at varying resolutions, and our European-ancestry discovery and metabolic-trait analyses motivate expanding multi-layer adipose QTL mapping to diverse ancestral populations. Third, while our cumulative posterior selection rule and sensitivity perturbations demonstrated high numerical stability across prior choices, colocalization and SMR provide model-based prioritization that now enables direct experimental perturbation—such as CRISPR base editing or isoform-specific knockdown—to establish causal mediation and therapeutic directionality^24,61^.

## Conclusions

By integrating five transcriptomic layers across subcutaneous and visceral adipose tissue, this study shifts the post-GWAS interpretation of T2D susceptibility loci from broad gene nomination to explicit RNA-feature hypotheses. Across 38 prioritized signal–gene relationships at 30 T2D signals, four non-gene-expression layers accounted for 87.0% of retained colocalizations and resolved regulatory relationships that were missed or misattributed by bulk gene expression alone. Combining locus-resolved candidate variants, external cohort corroboration, single-nucleus and cell-class genetic contexts, metabolic-trait profiles, and pharmacological annotations, our atlas defines whether follow-up assays should quantify total transcript abundance or specific exons, splice junctions, and transcript isoforms. This resource provides a practical, measurement-led blueprint for experimentally dissecting how non-coding genetic variation shapes adipose RNA regulation and contributes to T2D pathogenesis.

## List of abbreviations

2hGlu: 2-h glucose
ABF: approximate Bayes factor
ASPC: adipose stem and progenitor cell
BF: Bayes factor
BH: Benjamini–Hochberg
BMI: body mass index
CPM: counts per million
eQTL: expression quantitative trait locus
FG: fasting glucose
FI: fasting insulin
GE: gene expression
GWAS: genome-wide association study
HDL: high-density lipoprotein
HEIDI: heterogeneity in dependent instruments
HOMA-IR: homeostasis model assessment of insulin resistance
LD: linkage disequilibrium
LDL: low-density lipoprotein
MAF: minor-allele frequency
QTL: quantitative trait locus
SAT: subcutaneous adipose tissue
SD: standard deviation
SE: standard error
SMR: summary-data-based Mendelian randomization
SuSiE: sum of single effects
T2D: type 2 diabetes
TC: total cholesterol
TG: triglycerides
UMI: unique molecular identifier
VAT: visceral adipose tissue
WHR: waist-to-hip ratio
WHRadjBMI: waist-to-hip ratio adjusted for BMI.

## Declarations

### Ethics approval and consent to participate

This study analyzed publicly available genetic summary statistics, de-identified single-nucleus counts and accompanying metadata. No participants were recruited and no biological material was collected. Ethics approval and consent procedures for the contributing cohorts are described in their source publications.

### Availability of data and materials

Public inputs include the DIAMANTE fine-mapping archive, eQTL Catalogue release 7 (including GTEx liver, skeletal muscle, pancreas, whole blood and brain datasets), the METSIM adipose QTL release, GEO accession GSE176171 and the KOBS/RYSA cell-type eQTL release at Zenodo record 18880304. Metabolic-trait summary statistics were obtained from MAGIC (GWAS Catalog GCST90002227, GCST90002232, GCST90002238, GCST90002244 and GCST005179), GIANT–UK Biobank (Zenodo record 1251813), GLGC 2021, ADIPOGen (GCST001465) and the DIAGRAM 2018 BMI-unadjusted and BMI-adjusted releases. Islet eQTL data came from InsPIRE (Zenodo record 3408356), and UK Biobank LD matrices from the Broad Institute public release. Source URLs, releases and checksums are retained with the analysis tables. Summary results are provided in Supplementary Tables S1–S12, covering the discovery atlas, testability and sensitivity analyses, molecular features and conditional candidate sets, external adipose evidence, cellular expression and genetic contexts, metabolic-trait and tissue-context summaries, integrated follow-up readouts, tractability annotations and curated ChEMBL activity records. A permanent archive identifier for the final analysis and code release will be added before submission.

### Authors’ contributions

Yuelong Chen: Conceptualization, Methodology, Software, Validation, Formal analysis, Data curation, Visualization, Writing – original draft, Writing – review & editing. Shiya Huang: Data curation, Validation, Writing – review & editing. Tengxiao Lei: Data curation, Validation, Writing – review & editing. Yanci Wang: Data curation, Validation, Writing – review & editing. Haokun Han: Data curation, Validation, Writing – review & editing. Jiawei Li: Writing – review & editing. Lixin Cheng: Supervision, Writing – review & editing. Weidong Tang: Supervision. Jijun Tang: Supervision, Writing – review & editing. Nana Jin: Supervision, Writing – review & editing.

## Acknowledgements

We thank the participants and investigators of the source studies for making their data available for research.

